# Super-resolution microscopy of the ß-carboxysome reveals a homogenous matrix

**DOI:** 10.1101/086090

**Authors:** Matthew J. Niederhuber, Talley J. Lambert, Clarence Yapp, Pamela A. Silver, Jessica K. Polka

## Abstract

Carbon fixation in cyanobacteria makes a major contribution to the global carbon cycle. The cyanobacterial carboxysome is a proteinaceous microcompartment that protects and concentrates the carbon-fixing enzyme RuBisCO in a paracrystalline lattice, making it possible for these organisms to fix CO_2_ from the atmosphere. The protein responsible for the organization of this lattice in beta-type carboxysomes of the freshwater cyanobacterium *Synechococcus elongatus*, CcmM, occurs in two isoforms thought to localize differentially within the carboxysome matrix. Here we use widefield timelapse and 3D-structured illumination microscopy (3D-SIM) to study the recruitment and localization of these two isoforms. We demonstrate that this super-resolution technique is capable of successfully resolving the outer protein shell of the carboxysome from its internal cargo. We develop an automated analysis pipeline to analyze and quantify 3D-SIM images and generate a population level description of carboxysome shell protein, RuBisCO, and CcmM isoform localization. We find that both CcmM isoforms colocalize in space and time, prompting a revised model of the internal arrangement of the beta carboxysome.

## Introduction

RuBisCO (ribulose-1,5-bisphosphate carboxylase/oxygenase), the primary enzyme of carbon fixation, is ubiquitous in autotrophic organisms. However, it is notoriously inefficient and readily participates in energetically wasteful side-reactions with O_2_ (Andersson *et al*., 2008). Many organisms that rely on RuBisCO to acquire inorganic carbon tune the enzyme’s local microenvironment to achieve sufficient levels of carbon fixation. While plants can do this within membrane barriers, many single-celled organisms have evolved a strategy of proteinaceous compartmentalization.

The carboxysome is a bacterial microcompartment (BMC) found in cyanobacteria and prokaryotic chemoautotrophs. Composed of thousands of individual subunits, the carboxysome encapsulates RuBisCO and carbonic anhydrase (CA) within a selectively permeable icosahedral shell, making survival possible at atmospheric levels of CO_2_ (Figure 1A) (Rae *et al*., 2013; Kerfeld *et al*., 2016).

**Figure 1.**
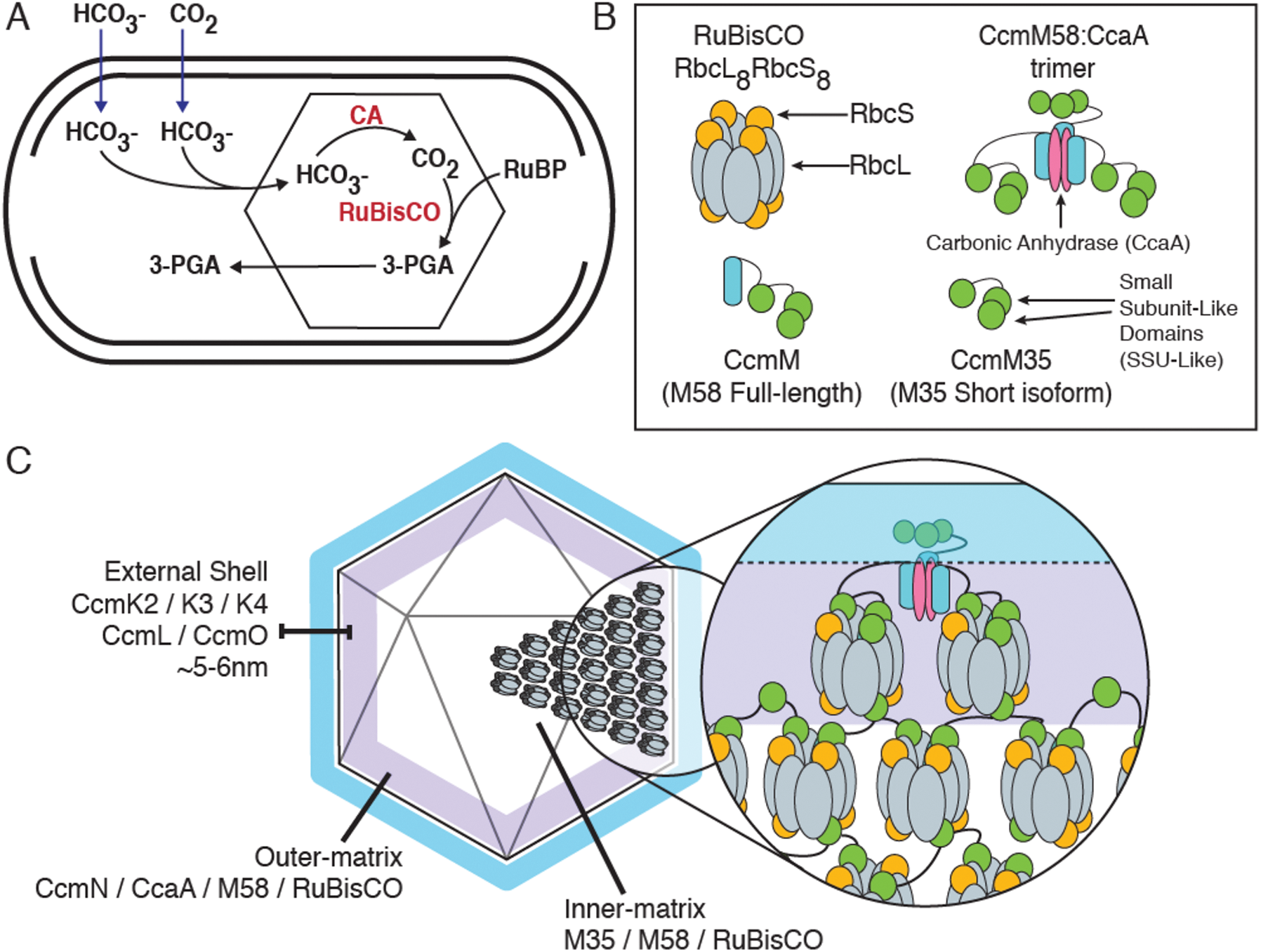
Current model of the cyanobacterial carboxysome internal structure. (A) *S. elongatus* utilizes the carboxysome to concentrate and protect its RuBisCO enzymes. Through a multi-step process, bicarbonate passes into the carboxysome, is converted to CO_2_ by carbonic anhydrase, and is then fixed by RuBisCO to form 3-phosphoglyceric acid (3-PGA). (B) Legend of proteins and protein complexes of the carboxysomes interior. *CcmM* is an essential carboxysome gene that is translated as two different isoforms, the full-length 58kDa M58 and a truncated 35kDa form, M35. CcmM has an N-terminal domain that complexes with CcaA carbonic anhydrase, and three repetitive C-terminal domains that have homology with the small subunit of RuBisCO. (C) These small subunit-like domains (SSU-like), which make up the majority of the M35 protein, are believed to displace a portion of the RuBisCO small subunits and physically tether the enzymes together within the core of the carboxysome, to the exclusion of the long M58 isoform which has been hypothesized to remain just beneath the carboxysome shell.

While evolutionarily distinct carboxysomes share many similar structural features, the RuBisCO in the beta-type carboxysome (ß-carboxysome) of the freshwater cyanobacterium *Synechococcus elongatus* PCC7942 is organized into a paracrystalline lattice (Kaneko *et al*., 2006). The structural protein CcmM has been implicated in connecting individual RuBisCO enzymes into an ordered matrix, resulting in the observed dense crystalline packing (Long *et al*., 2007; Cot *et al*., 2008, Long *et al*., 2010; Cameron *et al*., 2013). The full-length 58kDa CcmM protein of PCC7942 is characterized by an N-terminal carbonic anhydrase (CA) binding domain and three repeating domains toward the C-terminal end that have sequence homology to the small subunit of RuBisCO (Figure 1B) (Price *et al*., 1993). Due to an internal ribosomal entry site (IRES), CcmM is translated as two isoforms, a full-length protein (M58) and a 35kDa truncated version (M35) that consists of only the three small subunit like (SSU-like) domains (Long *et al*., 2007; Long *et al*., 2010).

The current model of ß-carboxysome internal structure suggests that the small subunit like domains (SSU-like) of CcmM displace some of the small subunits of the L8S8 RuBisCO holoenzyme and tether multiple enzymes together into a dense network (Figure 1C) (Long *et al*., 2007; Long *et al*., 2010; Rae *et al*., 2013; Kerfeld *et al*., 2016). Co-purification and yeast 2-hybrid experiments have shown that RuBisCO readily complexes with both isoforms of CcmM in PCC7942 (Long *et al*., 2007; Cot *et al*., 2008). In addition, fluorescence tagging experiments have demonstrated that M35 alone is capable of nucleating assembly of tagged RuBisCO if at least two of the SSU-like domains are present. However, M35 is not sufficient to cause encapsulation by recruiting shell components in the absence of M58 (Cameron *et al*., 2013). Densitometry analyses of western blot experiments have found imbalanced ratios of RbcS to RbcL (~5:8) in PCC7942 cell lysates, suggesting that the missing RbcS slots may be occupied by CcmM SSU domains (Long *et al*., 2011). These findings have supported a model of carboxysome architecture and step-wise assembly in which M35 nucleates a core of RuBisCO. In this model, M58 is limited to a layer beneath the shell where it can interact with CcmN, CA, and shell components. While this model predicts that the CcmM isoforms are differentially localized, these predictions have yet to be directly tested.

In order to test this model, we fluorescently tagged CcmM to visualize its assembly and the localization of CcmM isoforms *in vivo*. Previous work has shown that completely replacing CcmM with tagged fusions does not disrupt carboxysome function (Long *et al*., 2007; Cameron *et al*., 2015). Loss of either isoform of CcmM creates mutants that fail to produce carboxysomes and have a high CO_2_ requirement (HCR) for growth (Woodger *et al*., 2005; Long *et al*., 2010). However, complementation of such mutants with tagged copies of CcmM has been shown to rescue this phenotype (Long *et al*., 2007; Cameron *et al*., 2015).

Here we show the successful complementation of a *∆ccmM* mutant with a dually fluorescently tagged integration of *ccmM*. This dual tagging strategy does not significantly impair carboxysome formation, morphology, or function. In our timelapse fluorescence microscopy data we find no evidence of differential CcmM isoform assembly during carboxysome biogenesis. Using super-resolution 3D Structured Illumination Microscopy (3D-SIM), we examined the relative localization of carboxysome shell, CcmM, and RuBisCO proteins. Our 3D-SIM data does not support a model of carboxysome internal organization in which M58 is limited to a layer just beneath the external protein shell. These data support a revised model of carboxysome internal structure in which M35 and M58 isoforms are both integrated deep within the core of the carboxysome and are both involved with the early steps of carboxysome assembly.

## Results

### Fluorescent fusions to CcmM complement a deletion of the wildtype

We integrated an N- and C-terminal fluorescently-tagged copy of *ccmM* (*mTagBFP2-ccmM-mNeonGreen*) driven by the IPTG inducible promoter P*trc* into the genome of an *S. elongatus* PCC7942 ∆*ccmM*::*HygR* mutant strain (Figure 2A-B) and verified expression with western blot analysis (Figure 2C). This strategy allowed us to visually separate full-length CcmM from truncated M35 by blue fluorescence. We found that P*trc* was sufficiently leaky to produce protein without induction. Consequently we never induced the *∆ccmM* + *mTagBFP2-CcmM-mNeonGreen* strain, including for western blot analysis, unless otherwise noted. We blotted lysates of WT, ∆*ccmM*, and ∆*ccmM* + *mTagBFP2-CcmM-mNeonGreen* strains for the presence of CcmM (Figure 2C). In WT 7942 cells, we detected two bands at ~60 kDa and ~40 kDa, corresponding to the two native CcmM isoforms M58 and M35. In the *∆ccmM* strain, these bands are absent. In the *∆ccmM* + *mTagBFP2-CcmM-mNeonGreen* gene complement, we detected two bands at ~140 kDa and ~70 kDa, corresponding to molecular weight shifts of 60kDa (both fluorescent tags) and 30kDa (mNeonGreen alone), respectively. The observed weight shift of the tagged CcmM demonstrates that the CcmM IRES is not impaired by the presence of N- and C-terminal tags, and that both long and short CcmM isoforms are translated. We also made a reciprocally tagged version of the CcmM complement (mNeonGreen-CcmM-mTagBFP2), but did not use it for further analysis because western blotting showed the presence of protein degradation (Supplemental Figure S1).

**Figure 2.**
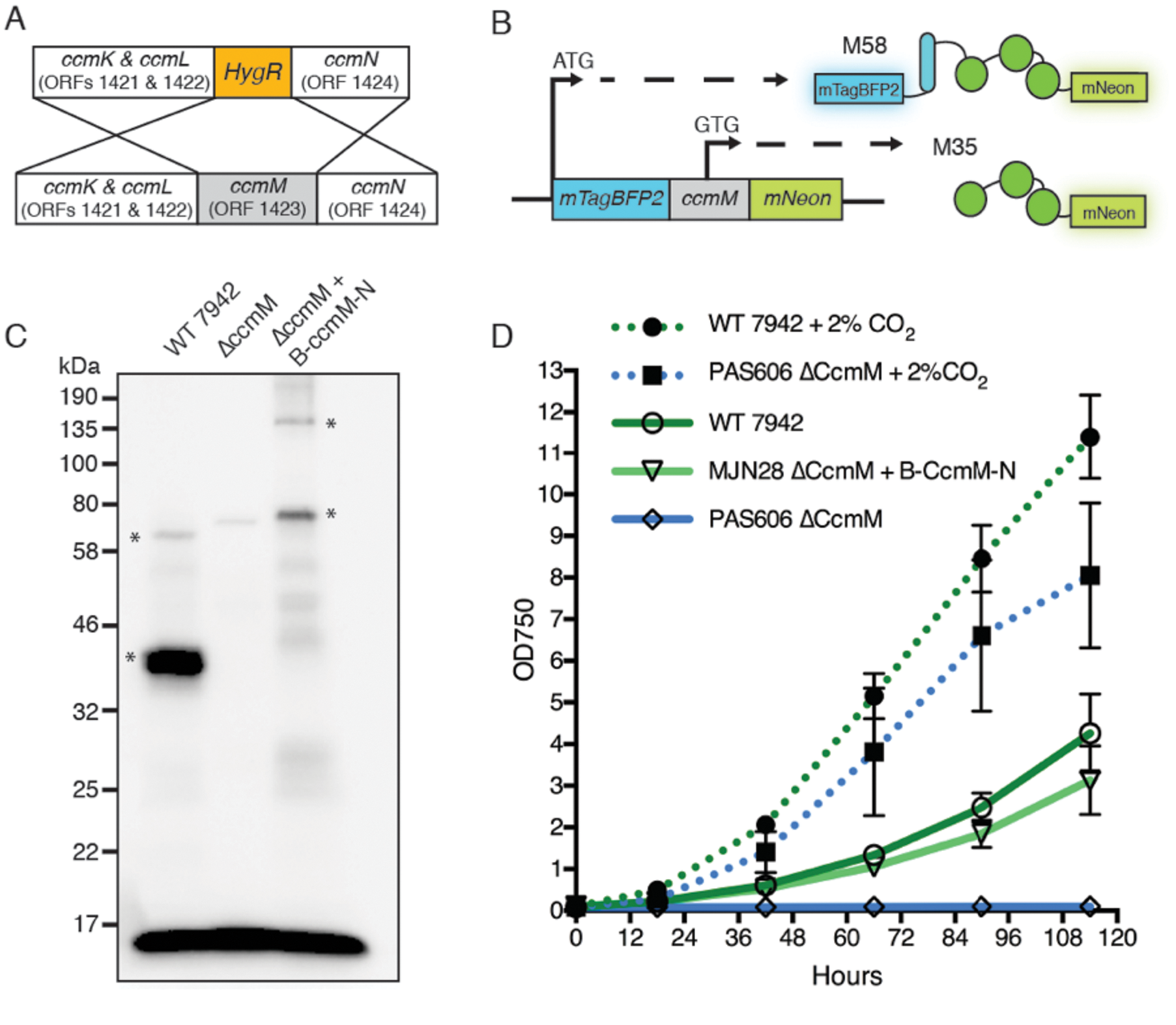
Fluorescently tagged *ccmM* integrated into the genome of a ∆*ccmM* knockout produces both CcmM isoforms and and restores growth in ambient air conditions. (A) The ∆*ccmM* knockout was made by integrating a hygromycin resistance marker into the genomic site of *ccmM*. (B) Diagram of fluorescently tagged *ccmM*. N- and C-terminal fusions of mTagBFP2 and mNeonGreen produces short and long isoforms with different fluorescent characteristics. This construct was integrated at Neutral Site II of the 7942 ∆*ccmM* genome to yield the ∆*ccmM* + *mTagBFP2-CcmM-mNeonGreen (+B-ccmM-N)* strain. (C) Anti-CcmM western blot of WT, ∆*ccmM*, and +*B-ccmM-N*. Full-length M58 isoform of CcmM is ~58 kDa, while the truncated M35 is ~35 kDa. The fluorescent fusions of mTagBFP2-M58-mNeonGreen and M35-mNeonGreen have expected molecular weights of approximately 113 kDa and 63 kDa respectively. (D) Growth rates of WT 7942 (n=6 ambient air, n=5 2% CO_2_), ∆*ccmM*, (n=6 ambient air and 2% CO_2_) and the +*B-ccmM-N* genetic complement (n=6 ambient air and 2% CO_2_). Dotted lines denote cultures grown with 2% supplemented CO_2_. Solid lines denote cultures grown in ambient air conditions.

Our tagged *ccmM* complement grew at a rate comparable to WT at atmospheric levels of CO_2_ (Figure 2D). We grew WT 7942, the *∆ccmM* mutant, and the ∆*ccmM* + *mTagBFP2-CcmM-mNeonGreen* complement in ambient air and 2% CO_2_ conditions, measured optical density at 750nm (OD_750_), and calculated doubling times between the 18- and 90-hour time points. Both WT and ∆*ccmM* cells cultured with 2% CO_2_ grew well with similar doubling times of 17.6 ± 0.4 hours (n = 5, Error = s.d.) and 15.8 ± 2.1 hours (*n* = 6, Error = s.d.) respectively. As expected, ∆*ccmM* mutant growth was dependent on supplemented CO_2_ (Long *et al*., 2010); these cells showed no growth in ambient air conditions. In contrast, the ∆*ccmM* + *mTagBFP2-CcmM-mNeonGreen* complement grew well in air with a doubling time of 23.3 ± 2.4 hours (*n* = 6, Error = s.d). This growth rate is similar to that of WT cells in air, which had a doubling time of 20.6 ± 1.8 hours (*n* = 6, Error = s.d.), indicating that our engineered strain is capable of fixing carbon, and that carboxysome function is not severely impaired by the dual-fusion *ccmM* complement. Because both the M58 and M35 isoforms of CcmM are required for growth without CO_2_ supplementation (Long *et al*., 2010), our data indicate that both isoforms produced by our dual fusion are functional.

Carboxysomes formed in the ∆*ccmM* + *mTagBFP2-CcmM-mNeonGreen* complement were structurally similar to those of WT cells. Using transmission electron microscopy, we imaged ultrathin sections of fixed cells from WT 7942, ∆*ccmM*, and ∆*ccmM* + *mTagBFP2-CcmM-mNeonGreen* strains (Figure 3). Carboxysomes in WT PCC7942 cells had an average maximum width of 179 ± 33 nm (*n* = 26, Error = s.d.). Carboxysomes in ∆*ccmM* + *mTagBFP2-CcmM-mNeonGreen* cells generally had a normal appearance and shape with visibly angled facets and often clear hexagonal geometry, but were noticeably larger, with an average maximum diameter of 319 ± 92 nm (*n* = 41, Error = s.d.). This is not unexpected; it has been previously reported that His-tagged *ccmM* complements increase carboxysome size (Long *et al*., 2007). Occasionally, dense masses at the cell poles were observed in ∆*ccmM* + *mTagBFP2-CcmM-mNeonGreen* micrographs, most likely indicating protein aggregation. Because we are using this strain without induction and no effective repressors exist, we cannot know if reducing expression would prevent aggregate formation. However, IPTG-induced cells observed with 3D-SIM display a dramatic increase in polar protein aggregation, often filling a large portion of the cell, confirming that overexpression of this construct leads to aggregation (Supplemental figure S2).

**Figure 3.**
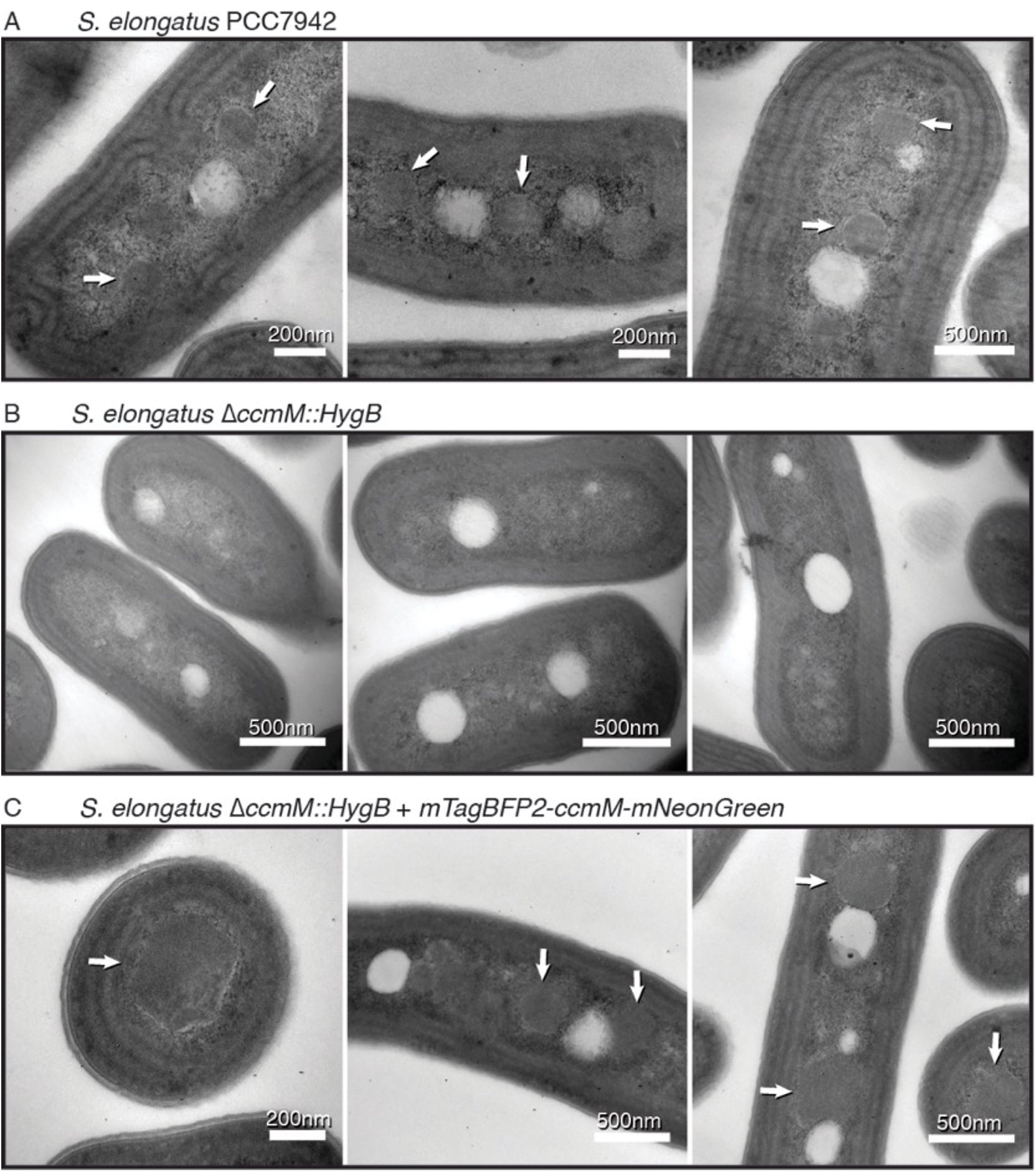
Transmission electron micrographs show the formation of carboxysomes in ∆*ccmM* cells complemented with fluorescently tagged *ccmM*. (A-D) Carboxysomes (white arrows) in wild type PCC7942 cells show characteristic hexagonal geometry. Diameters of WT carboxysomes labeled in micrographs range from 150 - 280 nm with an average of 179 ± 33nm (n = 26, Error = s.d.). (E-H) Micrographs of the ∆*ccmM* knockout show the absence of carboxysomes. (I-L) Integration of the fluorescently tagged *mTagBFP2-ccmM-mNeonGreen* construct into the into the ∆*ccmM* knockout rescues the formation of apparently normal carboxysomes. These carboxysomes had geometry similar to that observed in wild type cells. Carboxysomes with tagged CcmM appeared to range more widely in size than in wild type cells with an average diameter of 319 ± 92nm (n=41 Error = s.d.). Some (J & L) were of typical size (200nm in diameter), while others (K) are larger at 300-330 nm. Larger carboxysomes were also observed, like that seen in (I), which approaches 600nm at its widest point.

### Both CcmM isoforms share similar recruitment kinetics and localization

Both CcmM isoforms are recruited to growing carboxysomes with similar timing and kinetics. Using widefield microscopy, we imaged the CcmM complement (mTagBFP2-CcmM-mNeonGreen) at 10 minute intervals (Figure 4). Particle tracking revealed that both the CcmM isoforms assembled slowly, with an average assembly time of approximately 10.50 ± 5.39 hours (*n* = 8, Error = s.d.) for M58-mNeonGreen and M35-mNeonGreen, and 12.06 ± 7.27 hours (*n* = 8, Error = s.d.) for mTagBFP2-M58 (Figure 4C). This is longer than the ~6-7 hour maturation time previously reported for RuBisCO (Chen *et al*., 2013), perhaps as a result of temperature or light levels during imaging. Assembly times were estimated based on the time it took a particle to go from the intensity required for automatic particle detection (methods) to the first value ≥ 0.8 on the normalized intensity plot.

**Figure 4.**
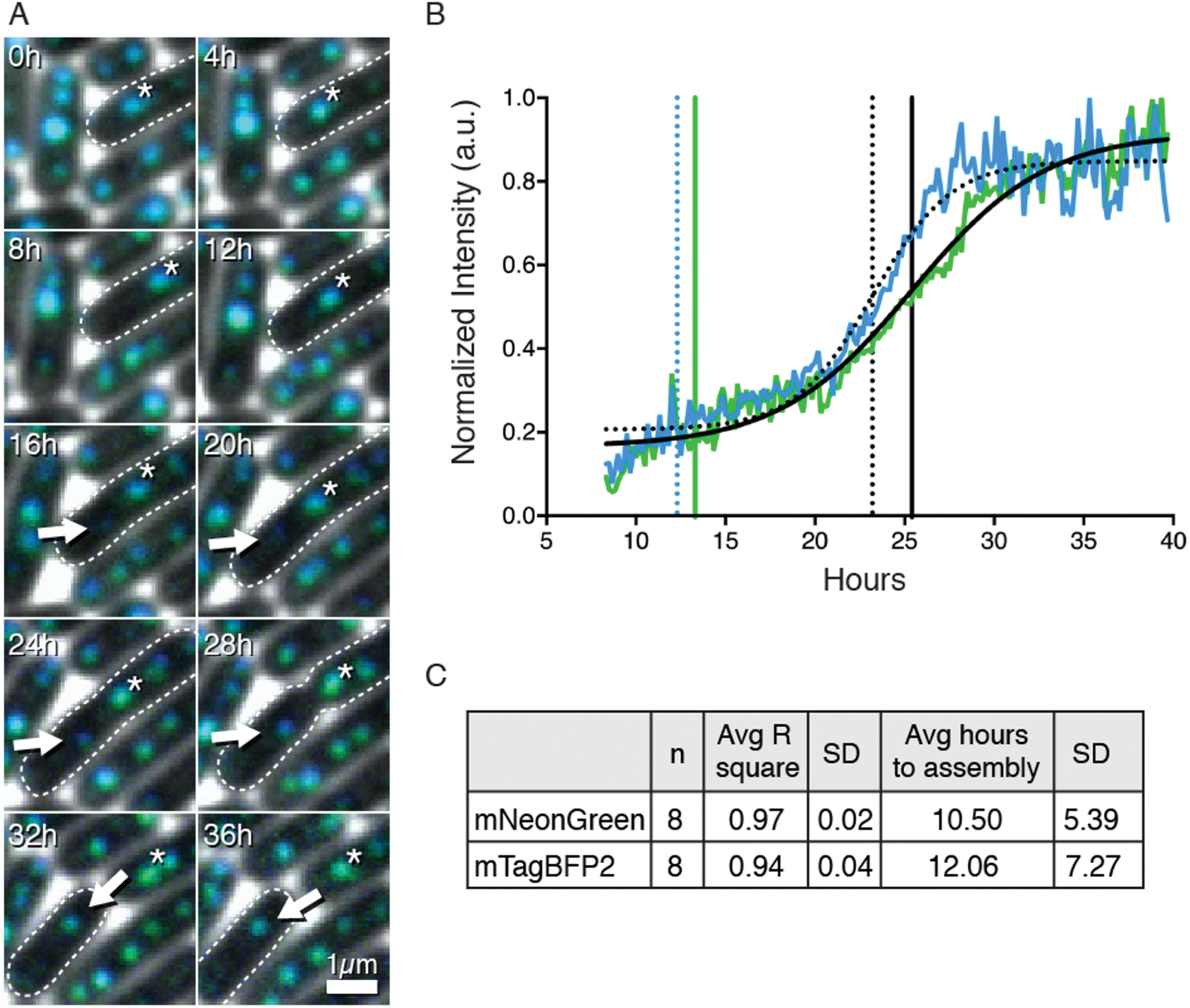
CcmM isoforms co-assemble at the site of new carboxysomes. (A) Merged montage of carboxysome biogenesis in the ∆*ccmM + mTagBFP2-ccmM-mNeonGreen* gene complement strain shows simultaneous co-localization of both blue (mTagBFP2-M58) and green (M58-mNeonGreen / M35-mNeonGreen) particles. White arrows indicate the new carboxysome. White asterisk indicates the parent carboxysome. (B) Channel specific intensity plots of carboxysome biogenesis depicted in panel A reveal sigmoidal-like dynamics in assembly timing in M35 and M58 CcmM isoforms. Vertical blue and green lines denote point at which automatic particle tracking first identified the new carboxysome. Tracks were extended backwards to measure background by manually measuring spot at or near the new particle (Methods). Four-parameter logistic equations were fit to each track; black solid or dotted vertical lines denote half-max values of each channel. (C) Statistics of sigmoidal curves fit to 8 measured biogenesis events find similar assembly dynamics in both green and blue channels. Assembly time was measured from first time point of automatic tracking to first value ≥ 0.8 on normalized intensity plot.

Next, a four-parameter logistic equation was used to approximately model the particle tracks. These models highlight similar assembly kinetics for both CcmM isoforms within the same carboxysome. Based on these fit curves, half-max values of green and blue particle assemblies were found to have an absolute average difference of 1.18 ± 0.68 hours (*n* = 8, Error = s.d.). If one CcmM isoform precedes the other during carboxysome biogenesis, the half-max values of each isoform would be expected to be significantly different. However, a comparison of half-max values from eight measured assemblies found no significant difference (Wilcoxon matched-pairs signed rank test, number of pairs = 8, two-tailed P = 0.945). In order to test if CcmM isoforms differ in rate of assembly, channel specific assembly times (average reported above) were analyzed and found not to be significantly different among paired fluorescent channels (Wilcoxon matched-pairs signed rank test, number of pairs = 8, two-tailed P = 0.375). This indicates that CcmM isoforms M35 and M58 assembled at relatively similar rates.

### M58 is not confined to a sub-shell layer in the carboxysome

Super-resolution microscopy of fluorescently tagged M58 and M35 show indistinguishable distributions in the carboxysome. We used 3D-Structured Illumination Microscopy (3D-SIM) to acquire super-resolution images of the ∆*ccmM* + *mTagBFP2-CcmM-mNeonGreen* strain, as well as fluorescently tagged RuBisCO (RbcL-sfGFP) and the shell protein CcmK4 (CcmK4-mTagBFP2 and CcmK4-sfGFP) in PCC7942. 3D-SIM images of strains with both GFP- and BFP-tagged CcmK4 resolved the external carboxysome shell as a ring structure in the xy-plane (Figure 5A-F), demonstrating that 3D-SIM provides sufficient resolution to identify peripherally localized proteins in the carboxysome. To confirm these rings were not an artifact of SIM reconstruction, we imaged a strain of PCC7942 integrated with a GFP labeled copy of the large subunit of RuBisCO (RbcL-sfGFP). This strain has been previously used successfully to study RuBisCO assembly and spacing (Savage *et al*., 2010; Chen *et al*., 2013). 3D-SIM of RbcL-sfGFP showed no ring-like structures, producing only solid regularly-spaced particles (Figure 5G-I). To test if M58 was in fact limited to a sub-shell layer, we imaged the *∆ccmM + mTagBFP2-CcmM-mNeonGreen* gene complement with 3D-SIM. Both CcmM isoforms produced solid objects similar to those observed with the RbcL-sfGFP strain (Figure 5J-N). These qualitative results suggested that the tagged M58 isoform of CcmM was not excluded from integration into the carboxysome core as would be expected if the protein was only present as a thin layer just beneath the exterior shell. In order to check that variations in signal intensity between experiments and cyanobacterial strains did not lead to poor signal-to-noise ratios and consequent artifacts, the reconstruction quality of each SIM data set was verified by the reconstructed intensity histogram (RIH) score from the SIMcheck ImageJ plugin (Ball *et al*., 2015). RIH scores for all conditions and data sets were of similar quality (Supplemental Table S1).

**Figure 5.**
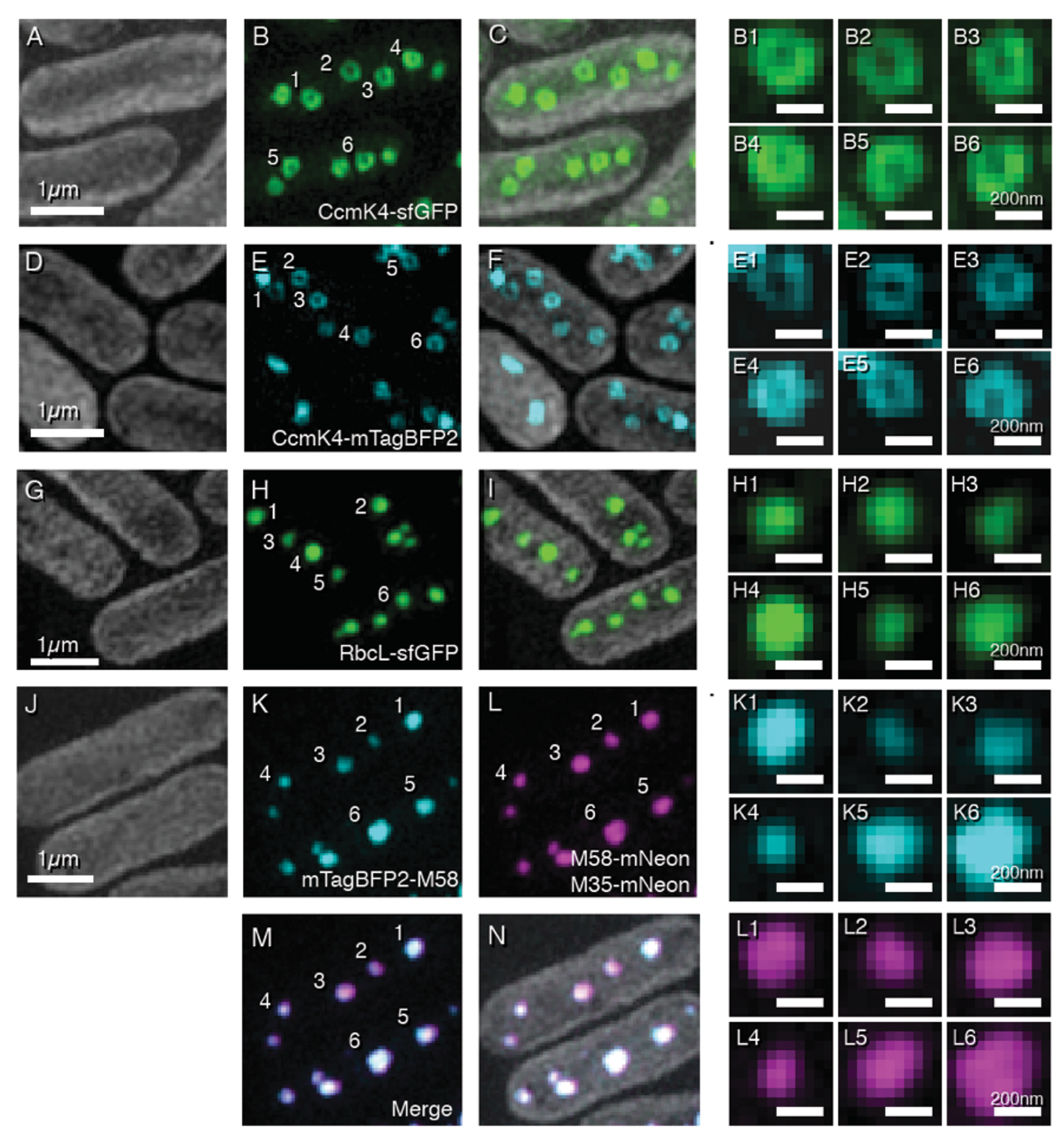
SIM of fluorescently tagged carboxysomes resolves the ring of the outer shell, and an absence of shell-like localization of either RuBisCO or CcmM. (A-C) SIM of CcmK4-mTagBFP2 resolves the carboxysome shell as a hollow ring, while (D-F) SIM of RbcL-sfGFP produces solid particles. (G-K) SIM of ∆*ccmM* + *mTagBFP2-ccmM-mNeonGreen* produces RuBisCO-like, apparently solid particles. M35/M58-mNeonGreen particles are pseudocolored magenta. Numbers identify individual carboxysomes and correspond to magnified panels on the right.

We developed an automated image analysis pipeline and a metric to quantify the degree to which fluorescently-labeled proteins were localized at the perimeter of the carboxysome versus uniformly distributed throughout the matrix. The max-to-center intensity (MTC) ratio is calculated for each particle as the peak intensity of the average radial profile divided by the intensity at the particle centroid. A perfectly uniform protein distribution would result in the peak particle intensity overlapping the particle centroid, and yield an MTC ratio of 1.00. Higher MTC values result from particles with decreased intensity at their centers, indicating a more ring-link appearance.

ANOVA testing confirmed that mean MTC values were significantly heterogeneous between all the labeled carboxysome proteins (Welch’s ANOVA, F_4,7378_=160.48, *P*<0.001). A Games-Howell test with *q*_critical_(k=5, df=∞, α=0.001)=5.484 was then used to make further pairwise comparisons of labeled carboxysome protein MTC values (Supplementary Table S2). This automated analysis of RbcL-sfGFP and CcmK4-sfGFP images confirmed our initial observations that carboxysomes appear fundamentally different between these strains (Games-Howell test, MD=0.02, *q*=22.26); similar results were obtained from the comparison of RbcL-sfGFP and CcmK4-mTagBFP2 (Games-Howell test, MD=0.03, *q*=26.38) (Figure 6A). RbcL-sfGFP particles had an average diameter of 244 ± 45 nm and an average MTC value of 1.002 ± 0.005 (*n* = 3505), with a maximum intensity ratio of 1.04. Based on this, we calculated an upper ratio threshold of 1.05, below which objects are likely to be uniform, by adding two standard deviations to the RbcL-sfGFP max ratio of 1.04. This threshold is denoted on MTC ratio versus diameter plots as a horizontal dotted line (Figure 6).

**Figure 6.**
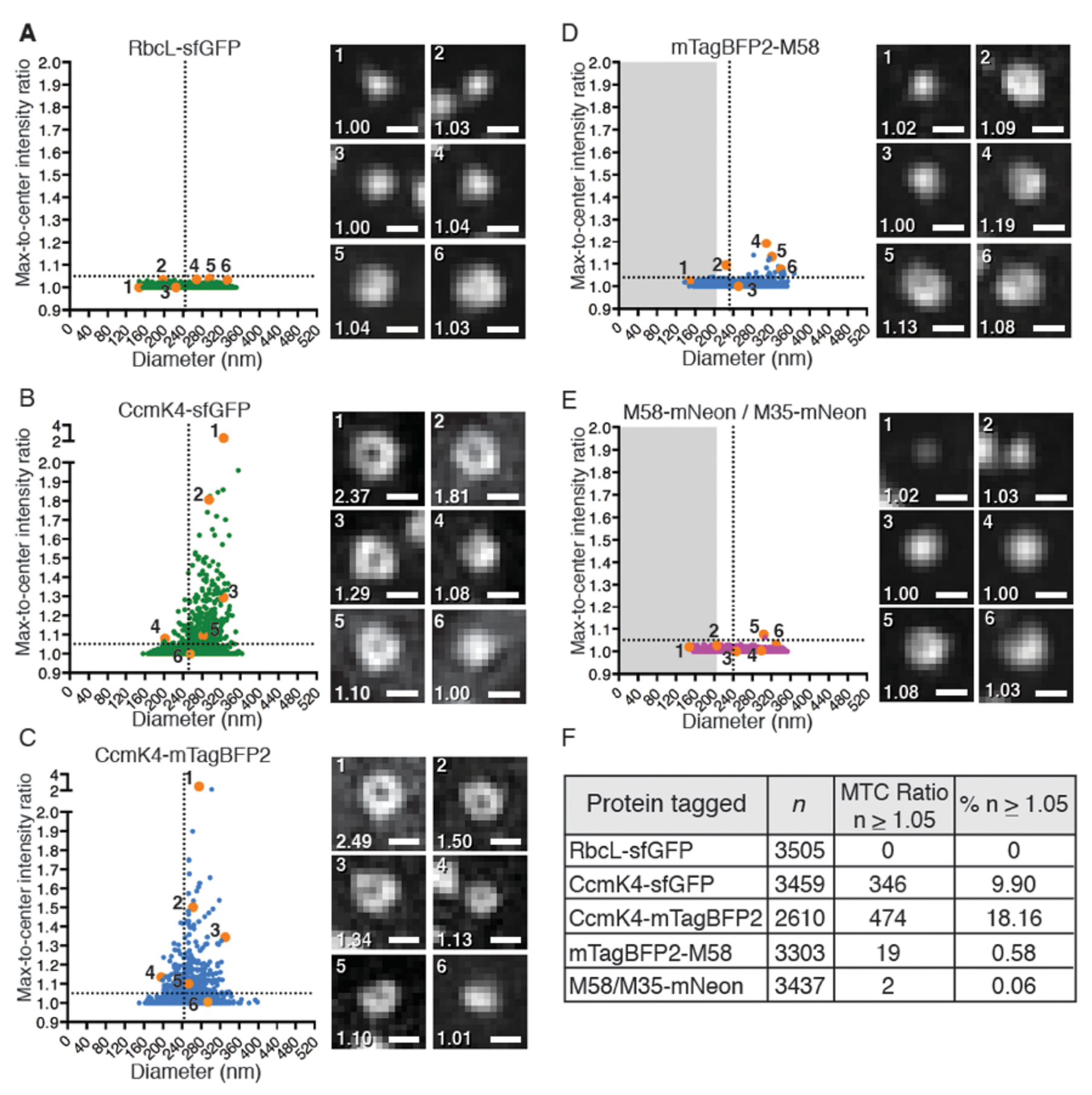
Analysis of 3D-SIM data. Using radially averaged intensity profiles of fluorescent particles in 3D-SIM data, maximum intensity to center (MTC) intensity ratios can measure the degree to which a fluorescent object is resolved as a ring. (A) Analysis of RbcL-sfGFP 3D-SIM data found no objects with a max-to-center ratio (MTC) greater than 1.04, with an average ratio of 1.002 ± 0.005 and average particle size of 244 ± 45 nm marked by the vertical line(error = s.d.). An upper ratio limit of 1.05, below which objects are likely to be solid, was calculated by adding two standard deviations to the RbcL-sfGFP max ratio. The 1.05 MTC cutoff is marked by a horizontal line on all plots. (B) Size versus intensity ratio plot of CcmK4-sfGFP shows a population of particles (345, 9.97%) with MTC ratios ≥ 1.05. (C) Plot of CcmK4-mTagBFP2 shows a two-fold increase in the fraction of ring-like particles compared to the GFP tagged version (474, 18.16%). (D) Analysis of particles in the blue channel of the ∆*ccmM* + *mTagBFP2-CcmM-mNeonGreen* strain, representing M58, finds only a few particles with noticeable ring-like appearance (19, 0.58%), with the vast majority of particles appearing solid. Based on the size of smallest detected ring in the tagged shell strains (196nm) and a shell thickness of 5-6nm, the minimum diameter at which ring-like shapes can be detected is calculated as 220nm; this is marked in gray. Ring-like particles within this region are unlikely to be resolved due to their small size. (E) Plot of particles in the green channel of ∆*ccmM* ± *mTagBFP2-CcmM-mNeonGreen* strain, representing M58-mNeonGreen and M35-mNeonGreen are predominantly solid with only 2 ring-like structures identified (2, 0.06%). (F) Table summarizing analysis statistics. In microscopy panels, scale bars are 200nm, and numbers denote MTC ratio.

In contrast to RbcL, we found that shell proteins displayed high MTC ratios. CcmK4-sfGFP particles had an average diameter of 253 ± 34 nm (*n* = 3459), and in comparison to RbcL-sfGFP particles, a significantly greater average MTC ratio of 1.023 ± 0.077 (Games-Howell test, MD=0.02, *q*=22.26, error = s.d.) (Figure 6H). Approximately 10% of CcmK4-sfGFP particles have MTC ratios ≥ 1.05, denoting a ring-like appearance (Figure 6B). Analysis of CcmK4-mTagBFP2 particles also revealed a significantly different mean MTC ratio compared to labeled RbcL-sfGFP (Games-Howell test, MD=0.03, *q*=26.38) (Figure 6H), with the CcmK4-mTagBFP2 particles having an average size of 244 ± 32 nm and average MTC ratio of 1.036 ± 0.093 (Figure 6C). Of note, the percent of ring-like particles identified in CcmK4-mTagBFP2 cells (18.16% ≥ MTC1.05) was significantly greater than the percentage found in CcmK4-sfGFP cells (9.90% ≥ MTC1.05) (Games-Howell test, MD=0.01, *q*=8.51) (Figure 6H). This increase in the fraction of rings identified with BFP labeled CcmK4 over those identified with GFP may be the result of the improvement in resolution due to the shorter wavelength of blue versus green fluorophores. Based on the smallest resolvable shell in the CcmK4-mTagBFP2 strain, which had an apparent diameter of 197nm, and accounting for an expected shell depth of 5-6nm, we do not expect to resolve by SIM even true ring-like particles smaller than approximately 220nm in diameter (Kaneko *et al*., 2006). This lower resolvable limit is denoted on plots of tagged M58 and of M35 particles by a grey region in Figure 6D-E.

Analysis of the ∆*ccmM* + *mTagBFP2-CcmM-mNeonGreen* gene complement identified a small number of tagged M58 particles as ring-like (Figure 6D). Our analysis found 0.58% (19 of 3303) of the mTagBFP2-M58 particles had MTC ratios of ≥ 1.05, with an average size of 233 ± 46 nm and average MTC ratio of 1.003 ± 0.010 (Figure 6F, error = s.d.). We found a significant difference in the MTC ratios of mTagBFP2-M58 particles vs both shell-labeled CcmK4-mTagBFP2 (Games-Howell test, MD=0.03, *q*=25.27) and CcmK4-sfGFP (Games-Howell test, MD=0.02, *q*=20.69) (Figure 6H). This supports qualitative observations that particles of the labeled M58 strain did not appear as ring-like as those in the strains with labeled carboxysome shells. mTagBFP2-M58 particles also had significantly different MTC ratios compared to RbcL-sfGFP particles (Games-Howell test, MD=0.001, *q*=10.05), but this difference was produced by only 19 ring-like particles in the mTagBFP2-M58 strain (Figure 6H). Taken together, these test results indicate that though mTagBFP2-M58 labeled carboxysomes have a small but statistically significant number of MTC ratios ≥ 1.05, the mean ratio is not similar to that produced by labeled carboxysome shell proteins, as would have been expected if the M58 isoform followed a strict sub-shell localization.

The *∆ccmM* + *mTagBFP2-CcmM-mNeonGreen* complement, which expresses both full-length M58-mNeonGreen and the short isoform M35-mNeonGreen, produced 2 particles with MTC ratios ≥ 1.05 (Figure 6E). Only 0.06% of particles were found above that threshold, with an average size of 244 ± 47 nm (n = 3437, error = s.d.) and average MTC ratio of 1.002 ± 0.005 (Figure 6F, error = s.d.). The mean MTC ratio of these particles was significantly different than those of both labeled CcmK4-sfGFP (Games-Howell test, MD=0.02, *q*=22.20) and CcmK4-mTagBFP2 (Games-Howell test, MD=0.03, *q*=26.34) (Figure 6H). Pair-wise comparison of M58-mNeonGreen and M35-mNeonGreen particles to RbcL-sfGFP found no significant difference in mean MTC ratios (Games-Howell test, MD=0.00005, *q*=0.57) (Figure 6G), indicating that the population of green fluorescent particles produced in the *∆ccmM* + mTagBFP2-CcmM-mNeonGreen strain were solid, similar to those observed in the strain with labeled RuBisCO.

## Discussion

### The carboxysome can support the addition of large foreign proteins via direct fusion to CcmM

The function of CcmM and other components of the carboxysome are robust to the addition of large protein fusions, further supporting these fusions as a means to localize foreign cargo to the carboxysome for engineering applications. Previous work has shown that tagged CcmM gene complements can rescue the HCR phenotype of CcmM mutants without significant impairment of carboxysome function compared to WT cells (Long *et al*., 2010; Cameron *et al*., 2015). We find that such complementation is also possible when fluorescent tags are fused to both ends of CcmM, significantly increasing the size of both the M58 and M35 isoforms. These results suggest that heterologous proteins could be simultaneously fused to multiple loci in the matrix, opening the door for localizing multipart enzymatic pathways inside the carboxysome.

### M35 and M58 assemble simultaneously during carboxysome biogenesis

M35 has been hypothesized to coordinate the organized paracrystalline cargo assembly of RuBisCO, with M58 limited to a peripheral sub-shell layer (Long *et al*., 2007; Cot *et al*., 2008; Long *et al*., 2010; Long *et al*., 2011; Rae *et al*., 2013). Current models of carboxysome biogenesis have proposed a primarily cargo-centric assembly pathway, in which RuBisCO and internal structural components form prior to shell encapsulation (Cameron *et al*., 2013; Chen *et al*., 2013). Based on FRAP imaging of carboxysomes that suggests matrix components are relatively static (Chen *et al*., 2013), we hypothesized that M35 would be present in pre-carboxysome assemblies prior to M58, with M58 appearing later during shell formation. However, we were unable to resolve any significant temporal difference between the CcmM isoforms. This concurrent timing indicates that M58 and M35 are present together even at the early stages of carboxysome assembly, with both isoforms involved in organization of the pre-carboxysome RuBisCO matrix. These findings are not necessarily in disagreement with current models of carboxysome assembly. Rather, they highlight how both CcmM isoforms are integrally involved in carboxysome biogenesis from the earliest stages. Assuming that the carboxysome core grows by addition of material to the outside of a stable paracrystalline RuBisCO assembly (Chen *et al*., 2013), these results also suggest that M58 does not assemble as a separate layer around a pre-existing M35/RuBisCO core.

### M58 is not exclusive to a subshell layer

3D-SIM experiments and quantitative analysis do not support a model of carboxysome internal structure in which M58 is limited to a sub-shell localization. The preexisting model described above proposes that M58 is not present within the primary M35-Rubisco matrix. This makes logical sense based on the fact that M35, which does not contain the N-terminal domain, presumably can achieve denser packing of RuBisCO enzymes.

We expected that if M58 was in fact limited to a sub-shell layer ~5-6nm beneath the outer surface, it would appear as a ring similar to CcmK4 by 3D-SIM. While SIM imaging of CcmK4 did reveal the expected ring-like distribution, carboxysomes with fluorescently-labeled M58 appeared predominantly as solid objects, with only a very a small population (0.58%) showing dips in fluorescence intensity at their centers. Consequently, we find no evidence by super-resolution microscopy to support a model of carboxysome structure in which M58 is excluded from the carboxysome core, suggesting that the internal carboxysome structure contains a mixed population of both M35 and M58 coordinating with RuBisCO. The finding that M58 is present throughout the carboxysome core raises questions about the compartment’s internal structure. For example, CcmM has been reported to interact with other carboxysome components (such as CcaA, CcmN, and CcmK) in yeast two hybrid and pulldown experiments (Long *et al*., 2007; Cot *et al*., 2008; Kinney *et al*., 2012), although the strength of these interactions is unknown. CcmN is essential for shell recruitment and assembly (Kinney *et al*., 2012), and thus is thought to localize at the carboxysome periphery (Rae *et al*., 2013). In addition, crystal structures of CcmM from *Thermosynechococcus elongatus* has revealed a homotrimer complex (Peña *et al*., 2010). Thus, CcmM may be involved in multiple structural and functional roles throughout the carboxysome (Figure 7).

**Figure 7.**
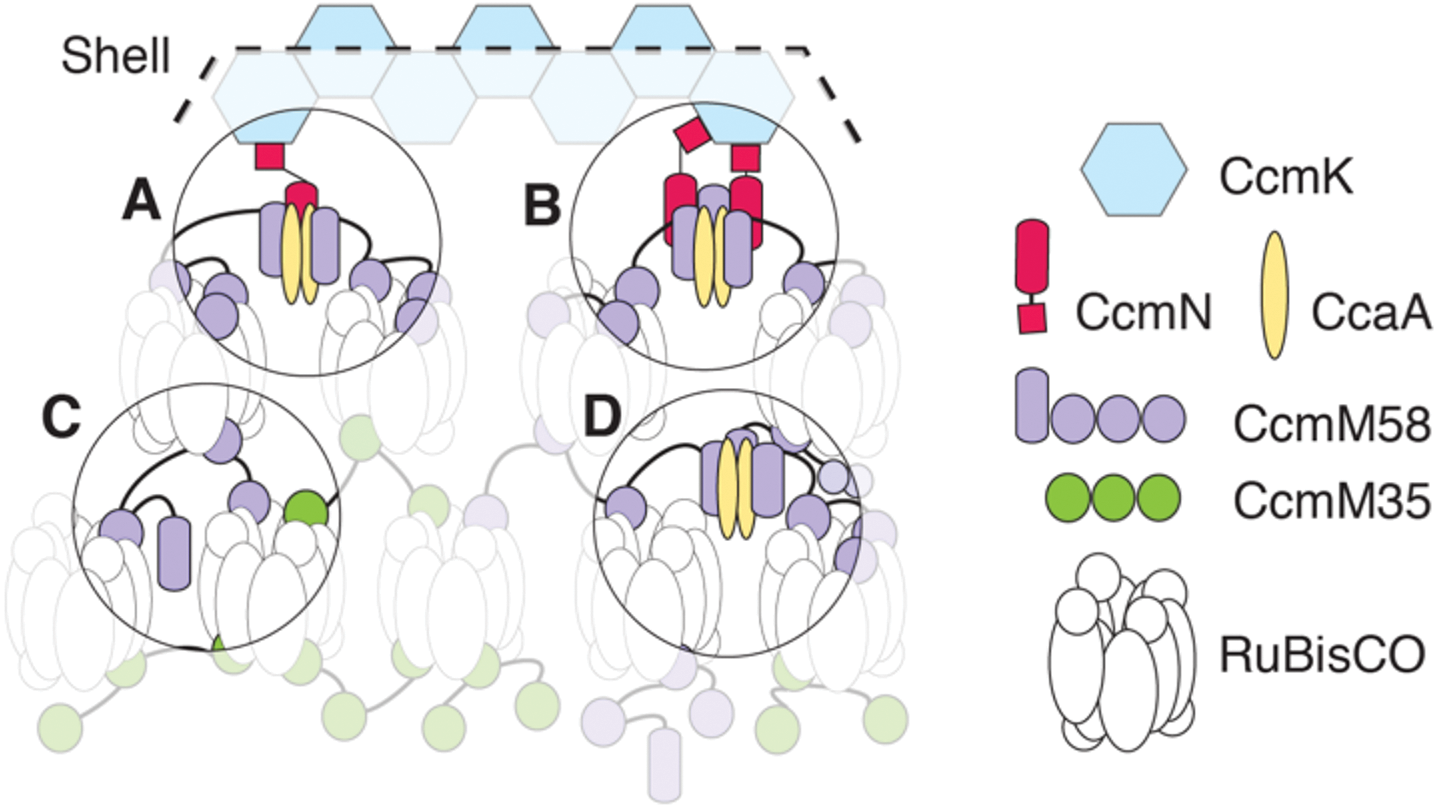
Mode of possible structural and functional roles of CcmM in the S. elongatus beta-carboxysome. (A-B) Full-length CcmM may form trimeric complexes through N-terminal interactions, which most likely include CcaA. Based on structural similarity between N-terminal domains of CcmN and CcmM, CcmN may be directly displacing units of CcmM in this complex or binding the timer to coordinate exterior shell recruitment (Kinney *et al*., 2012). (C-D) We find evidence supporting a model in which full-length CcmM58 is also present within the core of the carboxysome, which may involve single units of CcmM58 coordinating multiple RuBisCO enzymes as in (C) or may mean that CcmM58 is forming trimeric complexes within the interior.

Our findings also raise new questions about the structure and biogenesis of carboxysomes. For example, M58 is thought to play a role in recruitment of shell components (Cot *et al*., 2008; Long *et al*., 2011; Kinney *et al*., 2012; Cameron *et al*., 2013). However, shell proteins are recruited to carboxysomes only late in their maturation process (Chen *et al*., 2013). Therefore, if M58 is present even at the early stages of carboxysome assembly, other factors must be responsible for delaying shell recruitment. Furthermore, as our data indicate that CcmM58 is present deep within the carboxysome, it is possible that this protein is interacting with the carbonic anhydrase CcaA throughout the internal matrix of RuBisCO (Figure 7D). The nature of these factors is a promising avenue for future work.

## Materials and Methods

### Western blot analysis

For SDS-PAGE, log-phase (OD_750_ = 1.0) cultures of cyanobacterial strains were pelleted and frozen at -20°C. Frozen pellets were thawed in B-PER Bacterial Protein Extraction Reagent (ThermoScientific), without DNAse or Lysozyme, and normalized by wet pellet mass. Resuspended cells were mixed directly with SDS loading buffer and boiled. Boiled samples were run on a 4-20% Tris-Glycine NuPAGE gel (Invitrogen) at 120V. SDS-PAGE separated proteins were transferred to a nitrocellulose membrane with an iBlot2 transfer device (ThermoScientific).

The membrane was blocked with 5% skim milk in TBST (50mM Tris, 150mM NaCl, 0.1% Tween 20), then probed with 1:10,000 of anti-CcmM primary antibody (serum derived from rabbit, a gift of Ben Long), followed by a horseradish peroxidase-conjugated goat anti-rabbit secondary antibody (Agrisera AS09 602). The membrane was incubated with SuperSignal West Dura Extended Duration chemiluminescent substrate (Thermoscientific) and imaged with the Bio-Rad ChemiDoc system.

Western blotting for CcmM in WT, mutant, and transgenic cyanobacterial strain was replicated in two separate experiments, performed on different days, with independently collected culture samples (Supplemental figure S1).

### Transmission Electron Microscopy

For TEM experiments, 1mL of mid-log cultures were pelleted (6k rpm, 5 min) and concentrated 2x in sterile dH_2_O. The cells were then mixed 1:1 with fixative (1.25% formaldehyde, 2.5% glutaraldehyde, 0.03% picric acid, 0.1M sodium cacodylate buffer, pH 7.4), incubated at RT for 30 min, then pelleted and stored in fixative at 4°C overnight. Fixed cells were embedded in epon resin and cut into ultra-thin sections (~60-80nm). Sections were stained with uranyl acetate and lead citrate, then imaged on a Tecnai G2 Spirit Bio TWIN TEM.

### Cell culture

Wild-type, mutant, and modified strains of *Synechococcus elongatus* PCC7942 were grown in BG-11 media (sigma), with 1 g/L HEPES adjusted to pH 8.5. Cultures were grown at 30-35°C under constant light with an approximate intensity of 1000 lm/m^2^. Cultures were grown in 125ml or 250ml flat-bottom flasks, shaking at 150 rpm. Wild type 7942 and knockout complements were grown with ambient air conditions. Mutant *∆ccmM*::*HygR* cultures were supplemented with 2% CO_2_ and 25 µg/mL hygromycin B. Strains with fluorescently labeled carboxysome components, carrying kanamycin or spectinomycin resistance genes, were grown with 25 µg/mL kanamycin or 25µg/mL spectinomycin respectively. For the CcmM complement strain, *∆ccmM* + *mTagBFP2-CcmM-mNeonGreen*, cultures were grown with 5 µg/mL kanamycin and 25 µg/mL hygromycin B. For growth curve experiments, biological replicates were made by isolating colonies grown on BG11 agar with 1 g/L HEPES, 50mM HCO_3_^-^, and 1mM Na_2_S_2_O_3_. Two independent growth experiments were performed starting on different days, with 3 replicates in each experiment for a total of 6 biological replicates. One replicate of WT 7942 grown in 2% CO_2_ group was excluded due to significant evaporation in the culture. Isolates were screened by PCR to confirm the presence or absence of the WT *ccmM* gene. Growth curve cultures were prepared by washing starter cultures in fresh BG11 pH 8.5 media, then diluting to OD750 of ~0.1 in 40 mL BG11 in 250mL flat-bottom flasks.

### Image acquisition

For both conventional fluorescence and 3D-SIM experiments, 5µL of either mid-log cells (OD_750_ = 1-2) for timelapse experiments or early-log cells (OD_750_ = 0.5 - 1.2) for 3D-SIM were spotted on 2% agarose BG11 pads. Agarose pads were placed sample-side down in MatTek 35mm glass bottom microwell dishes with No. 1.5 coverslips. Sterile water was spotted within the dish to mitigate dehydration of the agarose pad, and the dish was sealed with parafilm. Though strains expressing labeled CcmM were under the control of the IPTG inducible promoter P*trc*, no induction was used with these cells due to adequate basal expression. Cells expressing RbcL-sfGFP were imaged on BG11 agarose pads with the addition of 50µM IPTG following previously reported protocols (Savage *et al*., 2010). SIM of uninduced RbcL-sfGFP cells also produced solid particles, but were not as bright as induced cells and produced several false-positives in our analysis pipeline (Supplemental figure S3). For timelapse experiments, samples were recovered for >1h at 34-35°C, under ~1000 lm/m^2^, and in ambient air, before being imaged using a Nikon TE2000 and a 100x oil objective. Cells were kept at 30°C under ~800 lm/m^2^ light intensity during timelapse, and imaged every 10min.

3D-SIM data was collected on a DeltaVision OMX V4 Blaze system (GE Healthcare) equipped with a 60x / 1.42 N.A. Plan Apo oil immersion objective lens (Olympus), and three Edge 5.5 sCMOS cameras (PCO). mTagBFP2 fluorescence was excited with a 405nm laser and collected with a 477/35 emission filter, mNeonGreen with a 514nm laser and 541/22 emission filter, and sfGFP with a 488nm laser and a 528/48 emission filter. Z-stacks of ~2 microns were acquired with a z-step of 125 nm and with 15 raw images per plane (five phases, three angles). Spherical aberration was minimized using immersion oil matching (Hiraoka *et al*., 1990). Super-resolution images were computationally reconstructed from the raw datasets with a channel-specific measured optical transfer function (OTF) and a Wiener filter constant of 0.001 using CUDA-accelerated 3D-SIM reconstruction code based on Gustafsson *et al. (*2008). Axial chromatic aberration was measured using TetraSpeck beads (Thermo Fisher) a nano-grid control slide and multi-channel datasets were registered using the image registration function in softWoRx. 3D-SIM data sets of each strain used for quantitative analysis were acquired in two independent experiments that performed with samples prepared on different days.

### Image analysis

SIM datasets were imported into MATLAB (Mathworks) using the bioformats reader, followed by thresholding and background subtraction. The threshold value was empirically determined and assessed by visualizing the resultant centroid position and mask in MATLAB and Imaris (Bitplane AG) respectively. Segmented objects were further analyzed if they met a cutoff for volume, surface area, sphericity, major-minor axis ratio, and mean radius.

We calculated the MTC (max to center) ratio for each object. From the centroid, the radially averaged intensity for each sub-sampled (0.5 pixels) radius was measured. This was done for each Z-plane and then averaged over the Z-axis. A polynomial curve was fit to the averaged radial profile of each segmented object in MATLAB. The most suitable polynomial fit (up to 6th order) was selected based on meeting the Bayesian Information Criterion. Then, the 1st and 2nd derivative of each fitted curve were calculated to find the root with the global maximum. The intensity at this root was calculated and divided by the intensity at the centroid. This ratio was calculated for each object and plotted versus its mean radius in MATLAB.

Notably, the resolvable carboxysome ring limit of 220nm that we estimate in our analysis is well above the ~110nm theoretical resolution limit of 3D-SIM (at 520nm wavelength; Gustafsson *et al*., 2000; Gustafsson *et al*., 2008). The difference is likely due to a combination of practical limitations along with the probable overestimation of carboxysome diameter in our analysis pipeline due to convolution of the true object size with the point spread function in the fluorescence images.

Timelapse data of ∆*ccmM + mTagBFP2-CcmM-mNeonGreen* were processed with ImageJ (FIJI) (Schindelin *et al*., 2012). Timelapse data from two sequential acquisitions of the same field of view were imported into FIJI with the Bio-Formats Import plugin with autoscaling, concatenated, and separated by channel. Channels were registered using rigid registration in the StackReg FIJI plugin, then background subtracted with a 50-pixel radius rolling ball. Particle tracking was performed in FIJI with the TrackMate plugin using the LoG detector (Laplacian of Gaussian filter) with 0.5µm estimated blob diameter, 0.5 threshold, and sub-pixel localization. Automatically generated tracks were manually extended backwards in time 20-30 frames by selecting 0.5µm diameter regions at or near site where the particle was first detected in order to capture a background fluorescence level. Particle tracks were corrected for photobleaching by dividing intensities by an average of 7 reference particles, normalized to the first frame of the time lapse, in each channel. Reference particles were objects observed to be relatively stable compared to particles that lost or gained fluorescence due to carboxysome biogenesis. The same reference particles were used for correction in both channels. To generate the timelapse montage, brightness and contrast values were manually adjusted separately in each channel to enhance visibility in the figure.

### Cloning and strain construction

Plasmids used for *S. elongatus* strain construction were made by Gibson assembly and sequence verified. Tagged CcmM plasmids were cloned by inserting the *ccmM* gene, along with mNeonGreen and mTagBFP2 coding sequences with additional flexible linker sequences, into the previously described pDFS21 neutral site II (NS2) integration plasmid downstream of the IPTG inducible P*trc* promoter (Savage *et al*., 2010). The *ccmM* gene was obtained directly from the *S. elongatus* PCC7942 genome via PCR with Phusion polymerase (NEB). The *ccmK4* gene for cloning shell fusion plasmids was obtained as a synthesized gBlock (IDT) and cloned into the pDFS21 integration plasmid (Supplemental Table S4). Annotated plasmid sequences have been 516 provided as supplemental materials.

*S. elongatus* strains were transformed with integration plasmids by incubating ~5 fold concentrated OD750 1-2 culture with 10-100ng plasmid at 30°C in the dark overnight, followed by plating transformants on selective BG11 agar (Clerico *et al*., 2007). The mutant ∆*ccmM* strain was made by integrating a hygromycin B resistance gene into the native *ccmM* ORF, followed by selective plating and screening (Sachdeva G., 2014).

#### Acknowledgements

We are grateful to Ben Long (ANU) for many helpful discussions, insightful comments on our BioRxiv preprint, and the kind gift of anti-CcmM serum. We thank Jennifer Waters and the Harvard Cell Biology Microscopy Facility for her guidance on SIM acquisition and analysis, Lin Shao for CUDA-accelerated SIM reconstruction code, Hunter Elliott (Harvard Medical School) for his great help and intellectual input on developing the SIM analysis pipeline, David Savage (UC Berkeley) for collaborating with us on exploring other microscopy methods, the Harvard Medical School EM facility for preparing samples and helping us with EM imaging. We would also like to acknowledge Gairik Sachdeva and Simon Kretschmer who originally constructed the ∆*ccmM::HygB* knockout strain in the Silver lab.

## Manuscript History

A previous version of this manuscript was posted to BioRxiv (https://doi.org/10.1101/086090).

## Supplemental Figures

**Figure S1.**
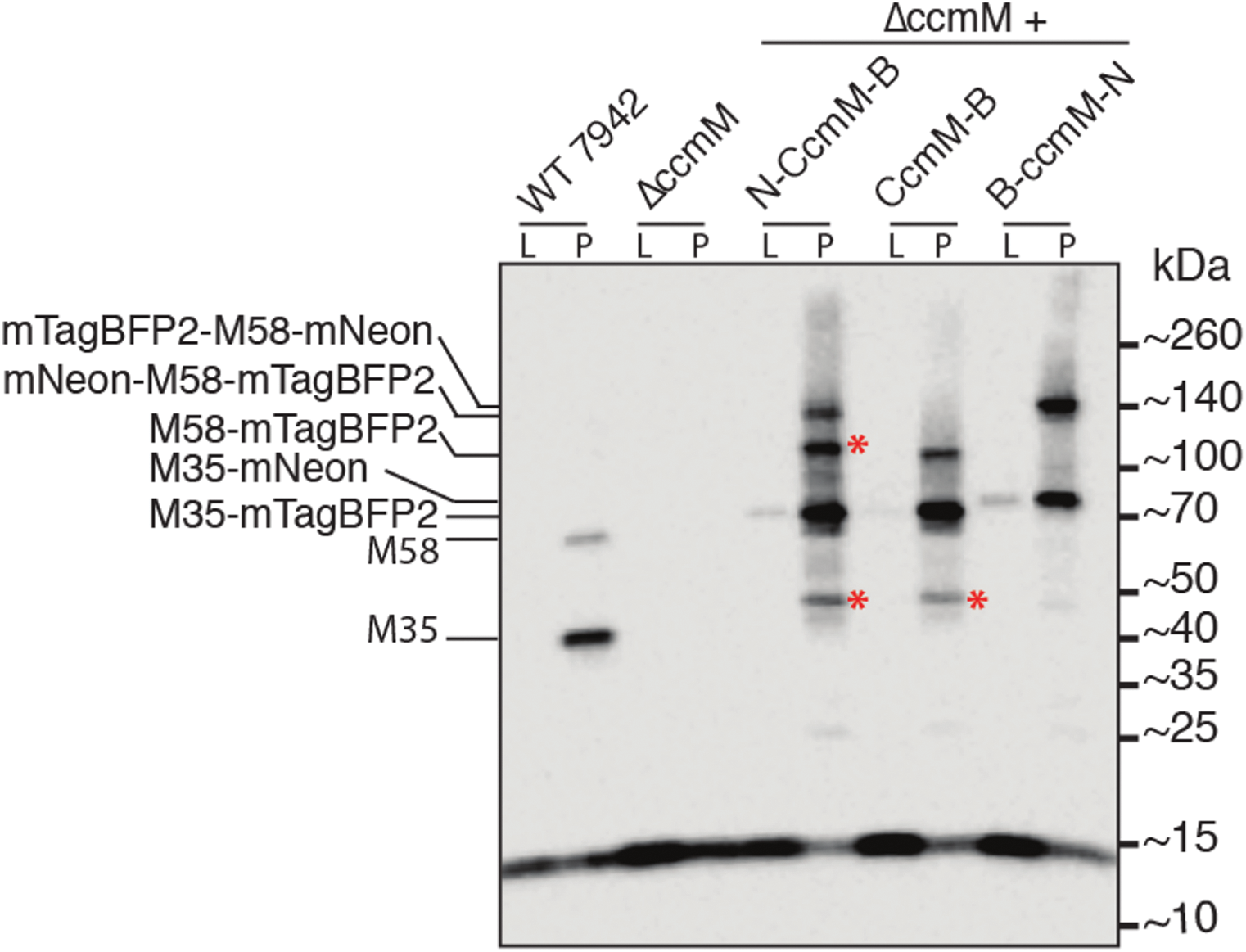
Protein degradation in variant ccmM gene complements. Western blot analysis of ∆*ccmM* + *mNeonGreen-ccmM-mTagBFP2*, and ∆*ccmM* + *ccmM-mTagBFP2* revealed prominent unexpected bands (marked by red asterisk) that could be indicative of protein degradation in the cell. The ∆*ccmM* + *mTagBFP2-ccmM-mNeonGreen* strain did not show similar unexpected bands. L = cell lysate and P = post-lysis pellet.

**Figure S2.**
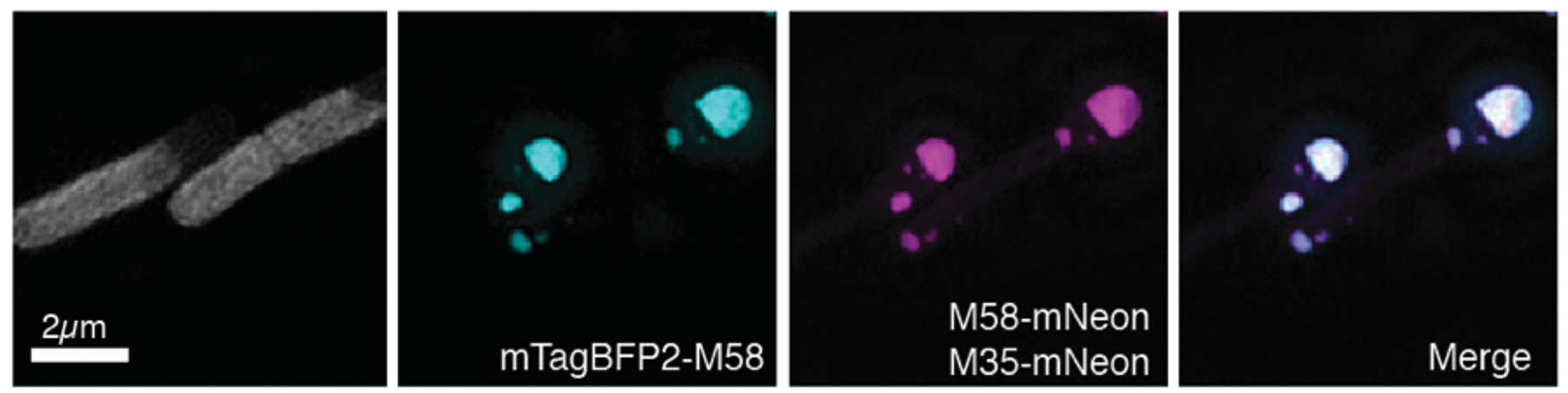
Induction of the Ptrc::∆*ccmM* + *mTagBFP2-ccmM-mNeonGreen* in S. elongatus leads to protein aggregation. SIM images of ∆*ccmM* + *mTagBFP2-ccmM-mNeonGreen* cells induced with 50µM IPTG show large aggregations of fluorescent signal near the cell polls.

**Figure S3.**
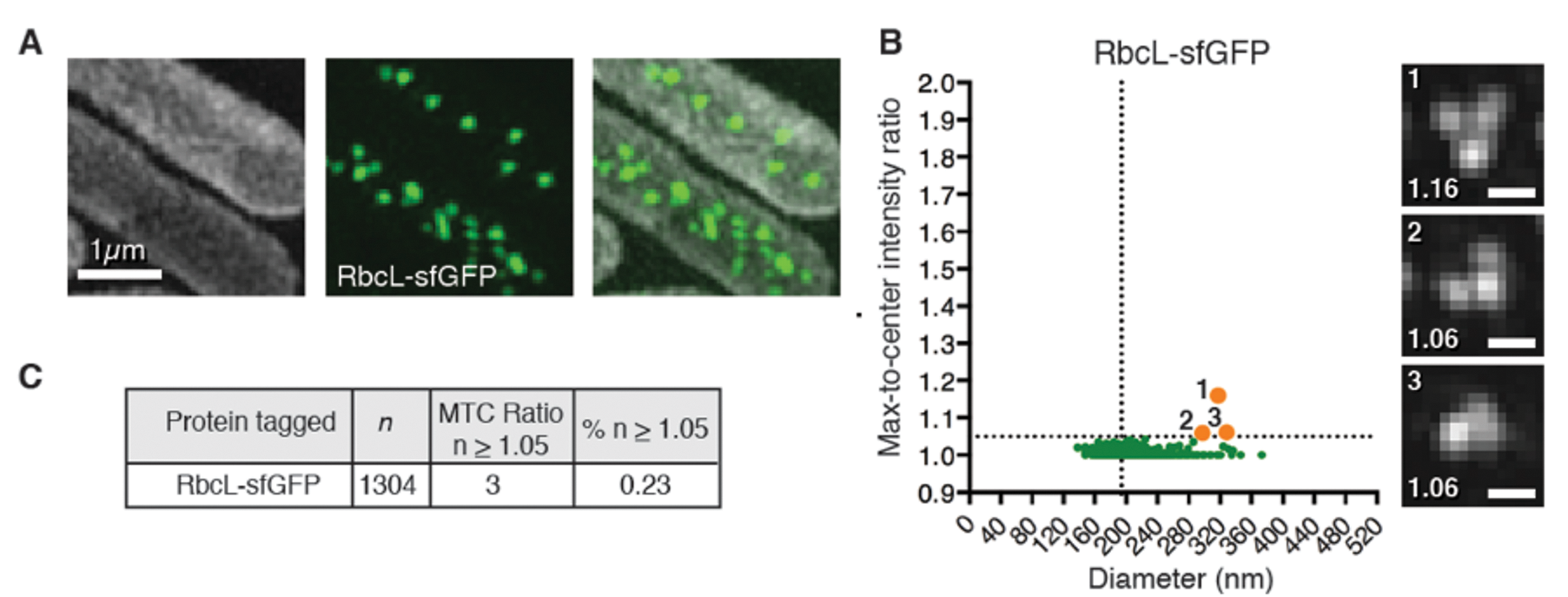
Uninduced cells expressing RbcL-sfGFP still produce solid particles. Qualitative data and image analysis results of PCC7942 RbcL-sfGFP cells without any IPTG induction show similar incidence of low MTC ratio solid particles compared to cells induced with 50µM IPTG. Three particles were identified as ring-like with MTC ratios > 1.05, but were confirmed to be three particle clusters incorrectly analyzed as single objects by our analysis program.

**Tables S1.** Structured Illumination Microscopy reconstruction scores. Reconstructed Intensity Histogram scores (RIH) were averaged across all image sets for each strain used for image analysis. Scores of 6-12 are considered good, and greater than 12 excellent. Columns with RIH scores are channel specific. Channel 608 captured the red auto-fluorescence of cells which was not used for any analysis. Error = Standard deviation.

| Strain | Tag | IPTG (μM) | # Imgs | Refractive Index | Wiener Filter | RIH (477) | RIH (528) | RIH (541) | RIH (608) |
| --- | --- | --- | --- | --- | --- | --- | --- | --- | --- |
| DFS724 | RbcL-sfGFP | 50 | 31 | 1.518 | 0.01 | | 12.74 ± 1.63 | | 2.78 ± 0.66 |
| AHC134 | CcmK4-sfGFP | - | 31 | 1.517 | 0.01 | | 13.15 ± 1.78 | | 3.53 ± 1.08 |
| MJN31 | CcmK4-mTagBFP2 | 0 | 29 | 1.517 | 0.01 | 8.28 ± 1.84 | | | 2.77 ± 0.96 |
| MJN28 | mTagBFP2-CcmM-mNeon | 0 | 47 | 1.517 | 0.01 | 9.45 ± 2.80 | | 12.58 ± 2.51 | 3.10 ± 0.84 |

**Table S2.** Statistical test results of particle MTC ratio comparison.

**A**
| Welch's ANOVA | Degrees of freedom | $F_s$ | $P$ |
| --- | --- | --- | --- |
| Among groups | 4.00 | 160.48 | 6.51 x 10<sup>-132</sup> |
| Within groups | 7377.44 |  |  |

| Games-Howell test | Mean difference | q | > $q_{crit}$ 5.484 ($\alpha=0.001$) |
| --- | --- | --- | --- |
| CcmM-mNeon vs CcmK4-sfGFP | 0.020465 | 22.20 | Significant |
| CcmM-mNeon vs CcmK4-mTagBFP2 | 0.033914 | 26.34 | Significant |
| CcmM-mNeon vs mTagBFP2-CcmM | 0.001283 | 9.56 | Significant |
| CcmM-mNeon vs RbcL-sfGFP | 0.000050 | 0.57 | Not significant |
| CcmK4-sfGFP vs CcmK4-mTagBFP2 | 0.013449 | 8.51 | Significant |
| CcmK4-sfGFP vs mTagBFP2-CcmM | 0.019182 | 20.69 | Significant |
| CcmK4-sfGFP vs RbcL-sfGFP | 0.020515 | 22.26 | Significant |
| CcmK4-mTagBFP2 vs mTagBFP2-CcmM | 0.032631 | 25.27 | Significant |
| CcmK4-mTagBFP2 vs RbcL-sfGFP | 0.033964 | 26.38 | Significant |
| mTagBFP2-CcmM vs RbcL-sfGFP | 0.001333 | 10.05 | Significant |

**Table S3.**
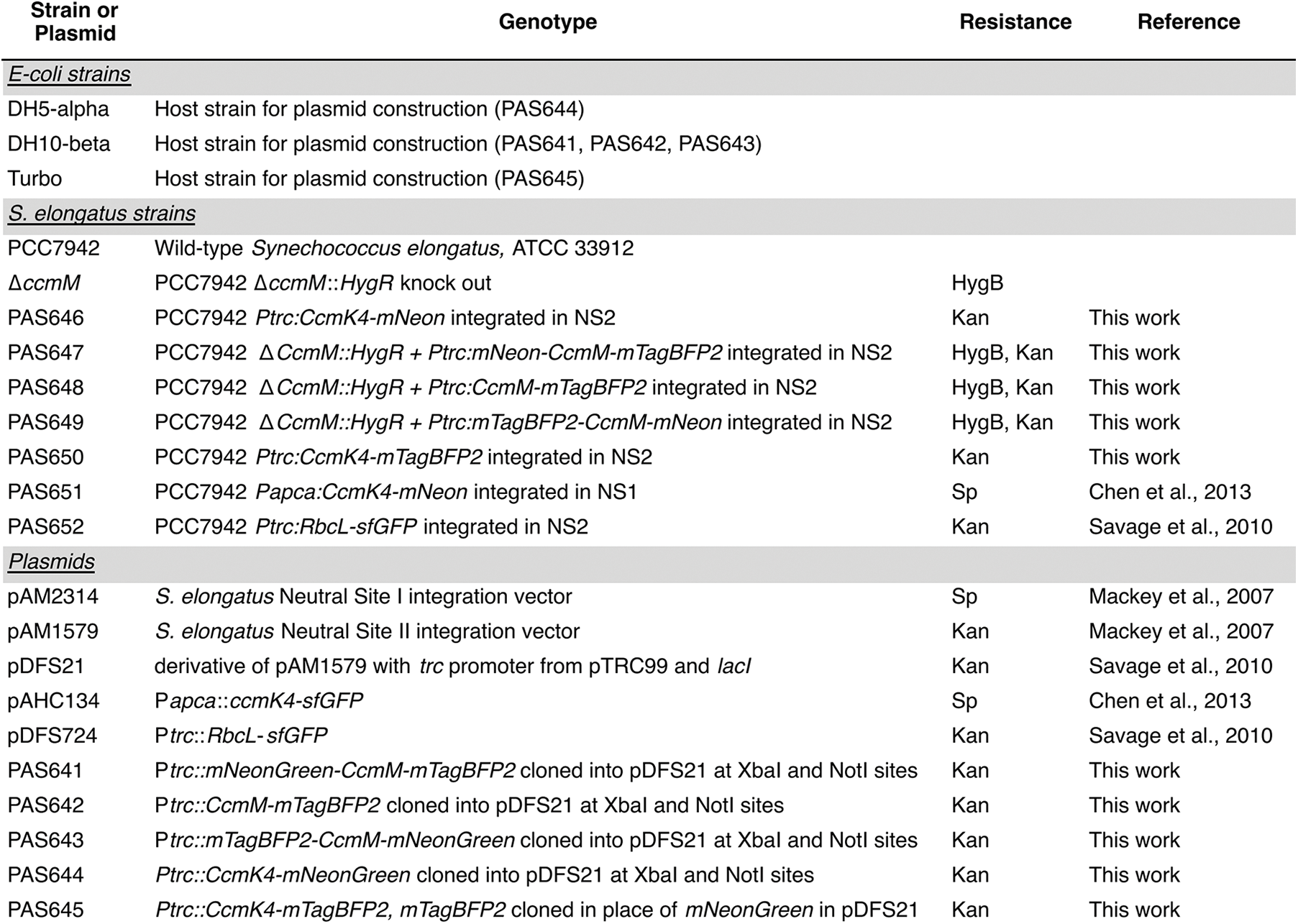
Strains and plasmids

**Table S4.**
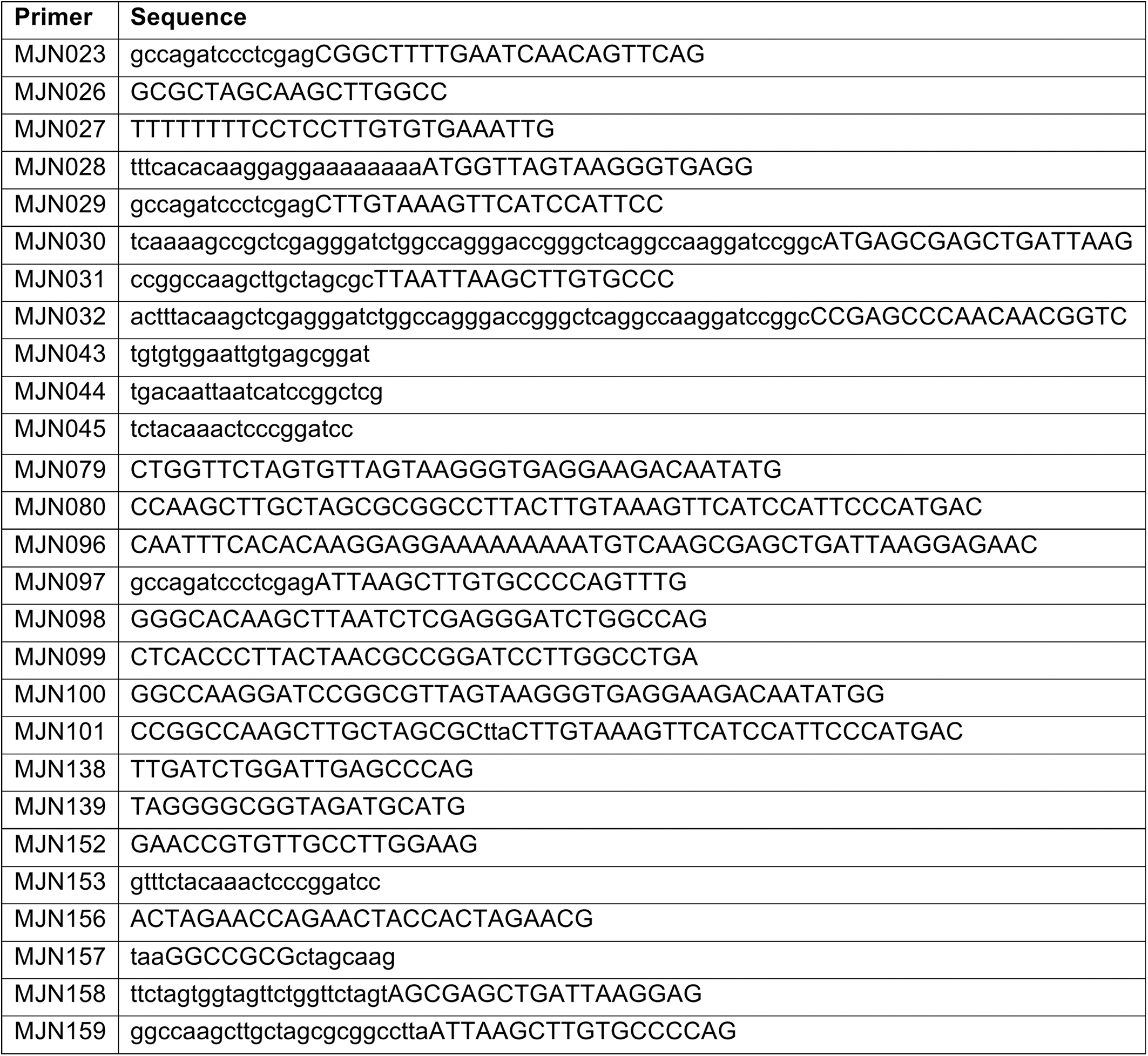
Primer sequences

## References

Andersson I, Backlund A. (2008). Structure and function of Rubisco. Plant Physio and Biochem. 46:275–291.

Badger MR, Hanson D, Price GD. (2002). Evolution and diversity of CO_2_ concentrating mechanisms in cyanobacteria. Funct. Plant Biol. 29: 161–173.

Badger MR, Price GD. (2003). CO2 concentrating mechanisms in cyanobacteria: molecular components, their diversity and evolution. J Exp. Bot. 54:609–622.

Ball G, Demmerle J, Kaufmann R, Davis I, Dobbie IM, Schermelleh L. (2015). SIMcheck: a Toolbox for Successful Super-resolution Structured Illumination Microscopy. Sci Rep. 5:15915.

Berla BM, Saha R, Immethun CM, Maranas CD, Moon TS, Pakrasi HB. (2013). Synthetic biology of cyanobacteria: unique challenges and opportunities. Front Microbiol. 4(246):1–14.

Cameron JC, Wilson SC, Bernstein SL, Kerfeld CA. (2013). Biogenesis of a bacterial organelle: the carboxysome assembly pathway. Cell. 155:1131–1140.

Chen AH, Robinson-Mosher A, Savage DF, Silver PA. Polka JK. (2013). The bacterial carbon-fixing organelle is formed by shell envelopment of preassembled cargo. PLOS One. 8(9):1–13.

Clerico EM, Ditty JL, Golden SS. (2007). Specialized techniques for site-directed mutagenesis in cyanobacteria. Methods Mol Biol. 362:155–171.

Cot SS, So AK, Espie GS. (2008). A multiprotein bicarbonate dehydration complex essential to carboxysome function in cyanobacteria. J Bacteriol. 190:936–945.

Gustafsson MGL. (2000). Surpassing the lateral resolution limit by a factor of two using structured illumination microscopy. J of Microsc. 198(2):82–87.

Gustafsson MGL, Shao L, Carlton PM, Wang CJR, Golubovskaya IN, Cande WZ, Agard DA, Sedat JW. (2008). Three-dimensional resolution doubling in wide-field fluorescence microscopy by structured illumination. Biophys J. 94:4957–4970.

Hiraoka Y, Sedat JW, Agard DA. (1990). Determination of three-dimensional imaging properties of a light microscope system. Partial confocal behavior in epifluorescence microscopy. Biophys J. 57(2):325–333.

Kaneko Y, Danev R, Nagayama K, Nakamoto H. (2006). Intact carboxysomes in cyanobacterial cell visualized by Hilbert Differential Contrast Transmission Electron Microscopy. J Bacteriol. 188(2):805–808.

Kerfeld CA, Melnicki MR. (2016). Assembly, function and evolution of cyanobacterial carboxysomes. 32:66–75.

Kinney JN, Salmeen A, Cai F, Kerfeld CA. (2012). Elucidating essential role of conserved carboxysomal protein CcmN reveals common feature of bacterial microcompartment assembly. 2012. J Biol Chem. 287(21):17729–17736.

Long BM, Murray BR, Whitney SM, Price GD. (2007). Analysis of carboxysomes from *Synechococcus* PCC7942 reveals multiple rubisco Complexes with carboxysomal proteins CcmM and CcaA. J Biol Chem. 282(40):29323–29335.

Long BM, Rae BD, Badger MR, Price GD. (2011). Over-expression of the ß-carboxysomal CcmM protein in *Synechococcus* PCC7942 reveals a tight co-regulation of carboxysomal carbonic anhydrase (CcaA) and M58 content. Photosynth Res. 109:33–45.

Long BM, Tucker L, Badger MR, Price GD. (2010). Functional cyanobacterial ß-carboxysomes have an absolute requirement for both long and short forms of the CcmM Protein. Plant Physiol. 153:285–293.

Peña KL, Castel SE, Araujo C, Espie GS, Kimber MS. (2010). Structural basis of the oxidative activation of the carboxysomal γ-carbonic anhydrase, CcmM. PNAS. 107(6):2455–2460.

Polka JK, Silver PA. (2013). Building synthetic cellular organization. Mol Biol Cell. 24:3585–3587

Price GD, Howitt SM, Harrison K, Badger MR. (1993). Analysis of a genomic DNA region from the cyanobacterium *Synechococcus* sp. Strain PCC7942 involved in carboxysome assembly and function. J Bacteriol. 175(10):2871–2879.

Price GD, Badger MR, Woodger FJ, Long BM. (2007). Advances in understanding the cyanobacterial CO_2_-concentrating-mechanism (CCM): functional components, Ci transporters, diversity, genetic regulation and prospects for engineering into plants. J Exper Bot. 59(7):1441–1461.

Rae BD, Long BM, Badger MR, Price GD. (2012). Structural determinants of the outershell of ß-carboxysomes in *Synechococcus elongatus* PCC7942: roles for CcmK2, K3-K4, CcmO, and CcmL. PLOS One. 7(8):1–11.

Rae BD, Long BM, Badger MR, Price GD. (2013). Functions, compositions, and evolutions of the two types of carboxysomes: polyhedral microcompartments that facilitate CO_2_ fixation in cyanobacteria and some proteobacteria. Microbiol Mol Biol Rev. 77(3):357–379.

Rae BD, Long BM, Whitehead LF, Förster B, Badger MR, Price GD. (2013). Cyanobacterial carboxysomes: microcompartments that facilitate CO_2_ fixation. J Mol Microbiol Biotechnol. 23:300–307.

Sachdeva G. (2014). Controlling fluxes for microbial metabolic engineering. Doctoral dissertation, Harvard University.

Savage DF, Afonso B, Chen AH, Silver PA. (2010). Spatially ordered dynamics of the bacterial carbon fixation machinery. Science. 327:1258–1261.

Schindelin J. et al. (2012). Fiji: an open-source platform for biological-image analysis. Nature Methods. 9(7): 676–682.

Schindelin J, Rueden CT, Hiner MC, Eliceiri KW. (2015). The ImageJ ecosystem: An open platform for biomedical image analysis. Mol Reprod Dev. 82(7-8):518–29.

Tanaka S, Sawaya M, Philips M, Yeates T. (2008). Insights from multiple structures of the shell proteins from the ß-carboxysome. Prot Sci. 18:108–120.

Woodger FJ, Badger MR, Price GD. (2005). Sensing of inorganic carbon limitation in *Synechococcus* PCC7942 is correlated with the size of the internal inorganic carbon pool and involves oxygen. Plant Physiol. 139(4):1959–1969.

Yokoo R, Hood RD, Savage DF. (2015). Live-cell imaging of cyanobacteria. Photosynth Res. 126(1):33–46.

